# Spatial and temporal variation in abundance of introduced African fig fly (*Zaprionus indianus*) (Diptera: Drosophilidae) in the eastern United States

**DOI:** 10.1101/2023.03.24.534156

**Authors:** Logan M. Rakes, Megan Delamont, Christine Cole, Jillian A. Yates, Lynsey Jo Blevins, Fatima Naureen Hassan, Alan O. Bergland, Priscilla A. Erickson

## Abstract

The African fig fly, *Zaprionus indianus* (Gupta), has spread globally from its native range in tropical Africa, becoming an invasive crop pest in select areas such as Brazil. *Z. indianus* was first reported in the United States in 2005 and has since been documented as far north as Canada. As a tropical species, *Z. indianus* is expected to have low cold tolerance, likely limiting its ability to persist at northern latitudes. In North America, the geographic regions where *Z. indianus* can thrive and seasonal fluctuations in its abundance are not well understood. The purpose of this study was to characterize the temporal and spatial variation in *Z. indianus* abundance to better understand its invasion of the eastern United States. We sampled drosophilid communities over the growing season at two orchards in Virginia from 2020-2022 and several locations along the East Coast during the fall of 2022. Virginia abundance curves showed similar seasonal dynamics across years with individuals first detected around July and becoming absent around December. Massachusetts was the northernmost population and no *Z. indianus* were detected in Maine. Variation in *Z. indianus* relative abundance was high between nearby orchards and across different fruits within orchards but was not correlated with latitude. Fitness of wild-caught females decreased later in the season and at higher latitudes. The patterns of *Z. indianus* abundance shown here demonstrate an apparent susceptibility to cold and highlight a need for systematic sampling to accurately characterize the range and spread of *Z. indianus*.

## Introduction

The African fig fly, *Zaprionus indianus* (Gupta), is an invasive drosophilid originating from tropical Africa (Yassin et al. 2008) that has spread to the Americas, Europe, and the Middle East in recent decades (Al-Jboory and Katbeh-Bader 2012, Kremmer et al. 2017, Molina-Rodríguez and Pérez-Guerrero 2019). Notably, *Z. indianus* was identified in Brazil in 1999 (Vilela 1999) where it has caused great losses as a pest of commercial fig crops (Oliveira et al. 2013). *Z. indianus* was first found in the United States in 2005 in Florida (van der Linde et al. 2006) and subsequently Virginia in 2012 (Pfeiffer et al. 2019). Populations in North America have been reported as far north as Minnesota in the United States (Holle et al. 2019) and Quebec, Canada (Renkema et al. 2013). Despite many incidental reports of its presence, no comprehensive studies have documented the geographical range or relative abundance of *Z. indianus* during a single growing season in the United States.

Despite the northward expansion of this species in North America, questions remain about the overwintering status of *Z. indianus* in these areas. *Z. indianus* likely does not survive winters in the north but rather disperses from southern refugia and recolonizes temperate habitats each year (Pfeiffer et al. 2019). Indeed, reports of *Z. indianus* in more northern states show inconsistent detections from year to year (Gleason et al. 2019, Holle et al. 2019), though (Joshi et al. 2014) speculated it overwintered in Pennsylvania. The exact locations where *Z. indianus* persists year-round remains to be investigated. However, while *Z. indianus* females can enter diapause and recover fertility afterwards (Lavagnino et al. 2020), males no longer produce progeny at temperatures lower than 15°C (Araripe et al. 2004), possibly limiting persistence at colder temperatures. Repeated sampling over a growing season is required to determine the timeline of its local colonization and extirpation in temperate environments.

*Z. indianus* is a generalist and uses multiple hosts in its original range in Africa (Yassin and David 2010) and both crops and native fruits where it has been introduced (Leão and Tidon 2004, van der Linde et al. 2006). In many of these hosts across the globe, *Z. indianus* are found at high numbers compared to other drosophilids (Silva et al. 2005, Roque et al. 2017, Pfeiffer et al. 2019). *Z. indianus* is primarily a secondary pest that largely infests damaged or decaying fruit, except in figs, but could become a pest of other crops (Bernardi et al. 2017, Pfeiffer et al. 2019). Assessing abundance across different fruits can delineate the habitats where *Z. indianus* can both exist and pose a threat.

The introduction of a new species such as *Z. indianus* can result in an upheaval of the native ecosystem, causing substantial biological and economic harm. Already established as a pest in Brazil, *Z. indianus* has the potential to become a significant pest in other areas. Characterizing an introduced species’ distribution is essential to understanding its impact and informing management solutions. For *Z. indianus*, however, the geographic areas and suitable habitats where it can become established are still not well understood. The purpose of this study was to characterize spatiotemporal variation in *Z. indianus* abundance to better understand its invasion of the eastern United States. To do so, we sampled drosophilid communities at two orchards in Virginia from 2020-2022 and several locations along the East Coast during the fall of 2022. Additionally, we investigated reproductive fitness of wild-caught females. We hypothesized that limited cold tolerance would result in reduced relative abundance of *Z. indianus* at more northern latitudes and that cold weather would correlate with reduced fitness and extirpation in temperate habitats.

## Materials and Methods

### Field Sampling

To assess seasonal abundance dynamics of *Z. indianus* populations, wild collections of drosophilids were conducted every 2-4 weeks in June through December from 2020-2022. Flies were collected from two orchards in Virginia (Charlottesville and Richmond, 116 km apart) that both harvest peaches (*Prunus persica*) in the summer and apples (*Malus domestica*) in the fall. Latitudinal sampling was conducted in 2022 between 30 September and 14 October. We collected flies from 11 sites in Maine, Massachusetts, Connecticut, Pennsylvania, Virginia, Georgia, and Florida. All sites were “pick-your-own” orchards except for Florida, which was a county park growing a wide variety of tropical fruits (but neither apples nor peaches were available). All sampling was conducted by random netting and aspiration of flies except for Charlottesville in 2021 and 2022, when 2L bottles baited with bananas (*Musa acuminata*) and baker’s yeast (*Saccharomyces cerevisiae*) were used as traps. *Z. indianus* were sorted from all other drosophilids and counted to determine relative abundance. We restricted our analysis of latitudinal variation to flies captured on apples for all sites except Florida. For Florida, we included all fruits in the analysis.

### Isofemale lines

Isofemale lines were generated from wild-caught flies collected in Connecticut, Pennsylvania, Georgia, and Florida. Additionally, isofemale lines were started from both Virginia orchards at two timepoints in 2022 (August and November). Following collection, all flies were held in bottles containing 50 mL cornmeal-molasses medium with yeast and a slice of banana for two days to encourage mating. Each female was placed in a vial with 10 mL cornmeal-molasses medium sprinkled with yeast. Females were incubated at 27°C and 50% relative humidity for one week. Isofemale line success was defined as the proportion of vials that successfully produced one or more larva.

### Analysis

Statistical analysis and plotting was performed with R (R Core Team 2021, v.4.1.2), *data*.*table* (Dowle and Srinivasan 2021), and *ggplot2* (Wickham 2016).

## Results

### Latitudinal Survey

We surveyed for *Z. indianus* along the east coast of the US between 30 September and 14 October during the 2022 season. We collected 6051 drosophilids (31.7% *Z. indianus*) from 11 orchards in seven states ranging from Florida to Maine. No *Z. indianus* were captured in the two orchards we sampled in Maine, and *Z. indianus* made up only 2% and 3% of drosophilids at the two Massachusetts orchards. The relative abundance of *Z. indianus* found on apples (or various hosts in Florida) varied widely across sites (Fig. 1; Table 1). In a linear regression (n = 11 orchards), latitude did not explain the variation in proportion of *Z. indianus* (P = 0.243, r = - 0.384). Female fitness, as measured by isofemale line success (n = 4 locations), decreased with increasing latitude but was not significantly correlated with latitude (linear regression: P = 0.103, r = -0.897, Table 1).

**Figure 1.**
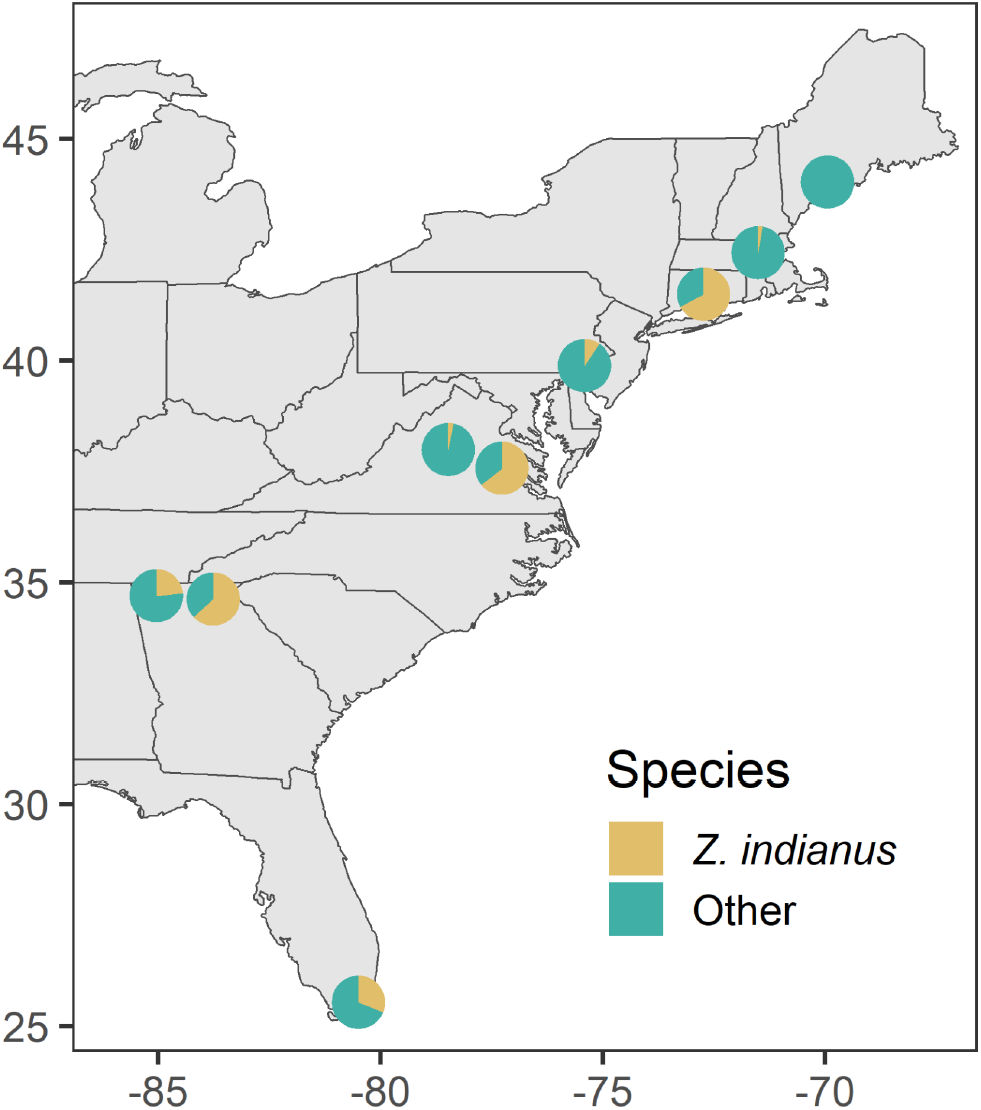
Abundance of *Zaprionus indianus* relative to other drosophilid species at selected sites sampled on the east coast of the US in early fall 2022. All individuals collected from apples except for Florida (various fruits). See Table 1 for additional information. The longitudes of Georgia pie charts were adjusted for visibility.

**Table 1.** Latitudinal variation in *Z. indianus* relative abundance in 2022. Isofemale line success refers to the proportion of females that produced offspring in the lab. The number of females tested is shown in parentheses.

| Location | Latitude | Longitude | Host | Relative abundance | Total collected | Isofemale line success | Collection Date |
| --- | --- | --- | --- | --- | --- | --- | --- |
| Maine | 44.025 | -69.943 | Apples | 0.0 | 20 | — | 8 Oct |
| Maine | 43.834 | -70.239 | Apples | 0.0 | 13 | — | 8 Oct |
| Massachusetts | 42.430 | -71.504 | Apples | 0.03 | 153 | — | 12 Oct |
| Massachusetts | 42.411 | -71.514 | Apples | 0.02 | 377 | — | 12 Oct |
| Connecticut | 41.494 | -72.730 | Apples | 0.67 | 446 | 0.55 (80) | 13 Oct |
| Pennsylvania | 39.885 | -75.410 | Apples | 0.10 | 208 | 0.66 (80) | 14 Oct |
| Charlottesville | 37.991 | -78.472 | Apples | 0.03 | 102 | — | 4 Oct |
| Richmond | 37.572 | -77.266 | Apples | 0.64 | 176 | — | 6 Oct |
| Georgia | 34.700 | -84.534 | Apples | 0.23 | 699 | — | 30 Sept |
| Georgia | 34.620 | -84.373 | Apples | 0.63 | 438 | 0.76 (80) | 30 Sept |
| Florida | 25.535 | -80.493 | Multiple | 0.31 | 2187 | 0.81 (200) | 2 Oct |

Within individual orchards, *Z. indianus* abundance varied on different fruits. At five out of seven timepoints with both apples and peaches available, *Z. indianus* was found at higher relative abundances on peaches than apples. Relative abundance was significantly higher on peaches when summed across all seven dates (chi-square: X^2^ = 268.41, df = 1, P < 0.001, Table 2). In Florida, we collected *Z. indianus* from eight different fruits. *Z. indianus* was most abundant on marula and least abundant on breadfruit, bael, and avocado (Table 3).

**Table 2.** Variation in *Z. indianus* relative abundance between apple and peach hosts. The total number of drosophilids collected is shown in parentheses.

| Location | Collection Date | Apples | Peaches |
| --- | --- | --- | --- |
| Charlottesville | 8/21/2020 | 0.19 (81) | 0.49 (300) |
| Charlottesville | 9/4/2020 | 0.39 (76) | 0.33 (662) |
| Charlottesville | 9/18/2020 | 0.18 (268) | 0.31 (416) |
| Charlottesville | 9/30/2020 | 0.18 (282) | 0.29 (92) |
| Richmond | 8/17/2022 | 0.83 (6) | 0.68 (433) |
| Pennsylvania | 10/14/2022 | 0.10 (208) | 0.54 (627) |
| Massachusetts | 10/12/2022 | 0.03 (153) | 0.11 (37) |
| Total |  | 0.16 (1074) | 0.45 (2567) |

**Table 3.**
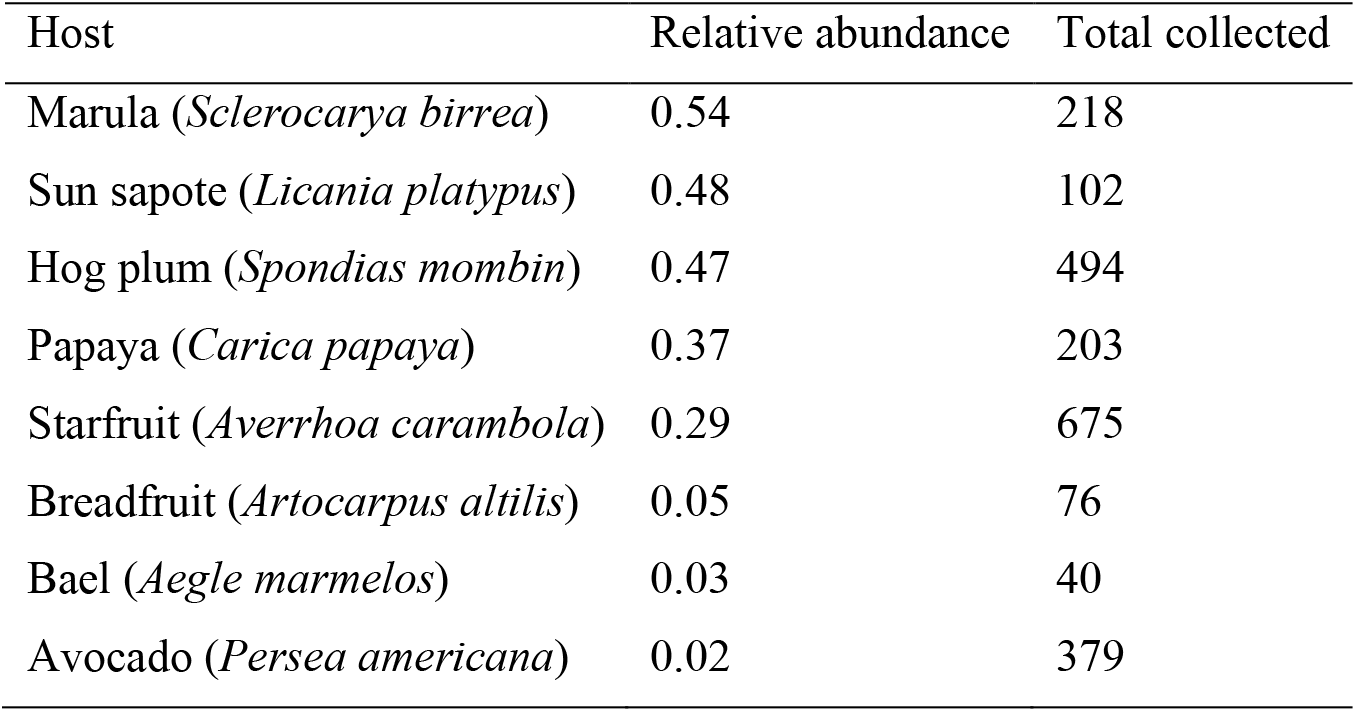
Variation in *Z. indianus* relative abundance across various fruits at a single park in Florida.

### Seasonal Abundance

From 2020 to 2022, we collected 24,648 drosophilids (27.1% *Z. indianus*) from orchards in Richmond, Virginia and Charlottesville, Virginia. *Z. indianus* abundance curves in both locations showed similar timing of population dynamics across years (Fig. 2). The first *Z. indianus* were generally captured around mid-July to early August, except in Charlottesville in 2022 when they were first captured in late June. The populations reached peak abundance around late August to early September. For most years, a second peak in abundance occurred later in the season, and numbers were low or undetectable by December. *Z. indianus* populations in Richmond reached higher relative abundance, peaking at ∼80-90% compared to a maximum of ∼40-45% in Charlottesville (Fig. 2). Female fitness was significantly higher in the early season for both Charlottesville (chi-square: X^2^ = 76.40, df = 1, P < 0.001) and Richmond (chi-square: X^2^ = 11.49, df = 1, P < 0.001, Table 4).

**Figure 2.**
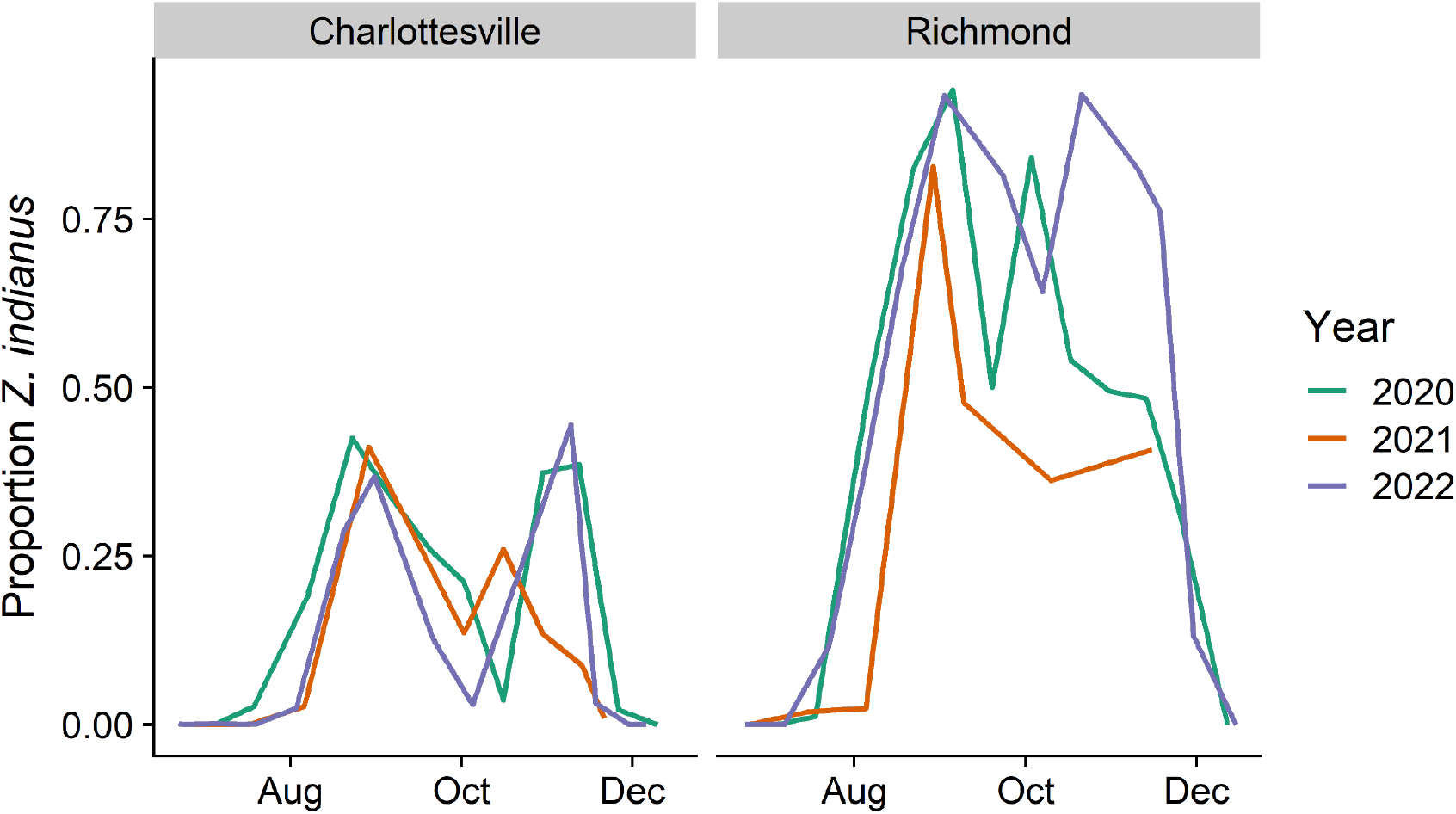
Seasonal abundance of *Z. indianus* collected from two locations in Virginia as a proportion of all drosophilids sampled. Flies were randomly collected from peaches and apples by netting and/or aspirating.

**Table 4.** Isofemale line success rate of wild-caught *Z. indianus* from two Virginia orchards in August and November 2022. The number of females tested is shown in parentheses.

| Orchard | August | November |
| --- | --- | --- |
| Charlottesville | 0.88 (117) | 0.29 (100) |
| Richmond | 0.84 (128) | 0.63 (100) |

## Discussion

We characterized variation in *Z. indianus* relative abundance in the eastern United States, establishing its distribution during a single season. Here, we report the first documentation of *Z. indianus* in Massachusetts, which was also our northernmost capture in 2022. In 2013, *Z. indianus* was reported at similar latitudes in Ontario, Canada in September and even further north in Quebec in October (Renkema et al. 2013). We detected no *Z. indianus* in our sampling of Maine orchards that occurred at similar latitudes as the reported Quebec captures. Overall captures in Maine were low, and *Z. indianus* could exist at undetectable levels in the population. Inconsistent detections from year to year have also been reported in Kansas (Gleason et al. 2019) and Minnesota (Holle et al. 2019). The lack of consistent detections across years suggests that *Z. indianus* does not survive year-round but rather disperses annually and sometimes does not colonize the same locations every year, especially at the edge of the range.

Across three years of sampling, typical first captures of *Z. indianus* in Virginia occurred in July or August despite detecting numerous other species in June and July. A similar trend was seen in Minnesota where *Z. indianus* was first captured even later in September and October (Holle et al. 2019). In Virginia, *Z. indianus* were largely undetectable by late fall even though other drosophilids were still captured. Late arrival and early decline compared to other species further supports yearly local extirpation and recolonization and differs from the seasonal dynamics of the invasive *D. suzukii*, which is thought to overwinter (Thistlewood et al. 2018). The consistent drop in abundance seen in mid-fall was notable and may be related to fruit preference and availability. Based on our field observations, peaches are the preferred food but rot faster than apples. During the period when most peaches have rotted but few apples are available, *Z. indianus* may lack suitable habitat and drop in abundance. Alternatively, another species may gain dominance, or a pathogen or parasite that affects *Z. indianus* may be common during this time. Dual peaks could also indicate bivoltinism; however, development takes 17.6 days at 22°C, which is much shorter than the time between abundance peaks (Nava et al. 2007).

The decrease in female fitness later in the season and with higher latitudes is consistent with our predictions. *Z. indianus* males reared at cool temperatures may take 9 days to produce offspring and may not recover fertility (Araripe et al. 2004). Whether the reduction in fecundity we observed is due to temperature effects on males, adaptive life history tradeoffs in females (as is seen in other drosophilids (Schmidt and Paaby 2008, Behrman et al. 2015)), or another cause remains a question. Assessment of fecundity in wild-caught and lab-reared flies could distinguish between phenotypic plasticity and adaptation for this trait.

Below an apparent threshold at Massachusetts/Maine, the effect of latitude on *Z. indianus* relative abundance was not straightforward. We saw large variation in relative abundance between nearby orchards at similar latitudes, including a nearly three-fold difference at Georgia orchards approximately 17 km apart, as well as within sites on different fruits. Differences in microhabitat and orchard management may play a larger role in determining relative abundance than latitude. Landscape cover, for example, has been shown to relate to *D. suzukii* abundance (Haro-Barchin et al. 2018). Our estimates of *Z. indianus* abundance are limited by uneven sampling efforts and methods across sites, and we were not able to sample further west in the species’ reported range. Additionally, by only sampling each latitudinal site once, our abundance estimates are likely influenced by weather conditions. Although we limited our latitudinal collections to a two-week period, the northern collection locales were more advanced in autumn seasonal phenology, so we cannot entirely deconvolute latitude and season in this study.

The patterns of *Z. indianus* distribution and seasonal dynamics we show here demonstrate an apparent susceptibility to cold as well as substantial variation in relative abundance at both large and small spatial scales. Our findings also highlight a need for systematic sampling to accurately characterize the range and spread of *Z. indianus* in real time and to track possible adaptive changes of this introduced species.

## Data Availability

All analysis scripts and raw data required to generate figures are available on Github: https://github.com/lmrakes/Zaprionus-field-collections-2022. Upon acceptance, data will also be deposited in Dryad.

## Acknowledgements

The authors would like to thank Shannon Jones and the students from the University of Richmond Integrated Science Experience program for assistance with early season collections. We would also like to thank Liam Miller, Cat Jalbert and Jerry He for assistance with fly collections and sorting. We are grateful to the owners and managers of all of the orchards we visited, especially Brian Campbell at Hanover Peach Orchard, Mechanicsville, VA, for making this study possible. This work was funded by NIH grant #1R15GM146208 to PAE, award #61-1673 from the Jane Coffin Childs Foundation to PAE, NSF BIO-DEB (EP): 2145688 to AOB, and startup funds from the University of Richmond. *Z. indianus* collections were conducted under permit #P526P-20-03908 from the USDA.

